# Topography and refractometry of sperm cells using SLIM

**DOI:** 10.1101/222000

**Authors:** Lina Liu, Mikhail E. Kandel, Marcello Rubessa, Sierra Schreiber, Mathew Wheeler, Gabriel Popescu

## Abstract

Characterization of spermatozoon viability is a common test in treating infertility. Recently, it has been shown that label-free, phase-sensitive imaging can provide a valuable alternative for this type of assay. Here, we employ spatial light interference microscopy (SLIM) to decouple the thickness and refractive index information of individual cells. This procedure was enabled by quantitative phase imaging cells on media of two different refractive indices and using a numerical tool to remove the curvature from the cell tails. This way, we achieved ensemble averaging of topography and refractometry of 100 cells in each of the two groups. The results show that the thickness profile of the cell tail goes down to 150 nm and the refractive index can reach values of 1.6 close to the head.

## 1 Introduction

Assisted reproduction is one of the major methodologies to treat human infertility. Human in vitro fertilization (IVF) was first introduced in 1978 to treat some forms of infertility. Presently, in Western societies, up to 4% of all children born are conceived using assisted reproduction technologies (ARTs)^1^. The Centers for Disease Control and Prevention (CDC) report from 2014 shows that 208,768 ART cycles were performed with 57,332 live births^2^. Several studies confirm the “male side” is found to be responsible for 20-30% of infertility cases and contribute to 50% cases overall^3^. The factors may be low sperm concentration, poor sperm motility, or abnormal morphology and these can happen singularly or in combination^4^. Other problems are hormone imbalances or blockages in the male reproductive organs. In about 50% of cases, the cause of male infertility cannot be determined^5^. Current research is focused on developing a non-invasive method to evaluate the semen morphology, with the possibility to use the sperm after the evaluation. In this scenario a method to evaluate the sperm parameters can be a very useful tool to increase the birthrate success after ARTs.

Here we show that SLIM^6–8^, a highly sensitive quantitative phase imaging (QPI) method, can be used to study unlabeled sperm cells in great detail. QPI is an emerging method that provides quantitative information about transparent specimens, with nanoscale sensitivity and in label-free mode^9^. Various implementations of QPI have been used recently to study the topography of gametes, assuming a certain refractive index^10–16^. Here, for the first-time to our knowledge, we measure independently the thickness and refractive index of the cells. The results for topography and refractometry of the tail are in the expected range, as revealed by electron microscopy^17^. Remarkably, we found that the refractive index of the cell mid-piece (neck) is very high. It is known that this region is mitochondria-rich, which may explain this interesting finding^18^.

## 2 Materials and Methods

### 2.1 Sperm Cell Preparation

Straws of frozen bovine semen from bulls previously tested for in vitro fertilization (IVF) were thawed at 37 °C for 40 sec. The sample was selected by centrifugation (25 min at 300×g) on a discontinuous gradient (45–80% in Tyrode’s modified medium without glucose and Albumin). The pellet was reconstituted into 2 ml of IVF medium and centrifuged twice, at 160 and 108×g for 10 min. For each sample pellet was diluted with IVF medium at the concentration of 1×10^6^ sperm/ml^19^. The final sample was divided into sample of dry sperm and sample in PBS (Phosphate buffer saline).

#### Dry samples

10 microliters of selected sperm were smeared onto a clean microscope glass slide (24-50 mm), left to dry for 10 minutes and finally fixed in 98% ethanol for 10 minutes^20^. Slides were stored at 4°C until the day of the evaluation.

#### Sample in suspension

10 microliters were mixed with 50 microliters of PBS and smeared onto a clean microscope glass slide. To avoid the liquid evaporation samples were overlaid with a cover slide and sealed with nail polish. Slides were stored at 4°C until the day of the evaluation.

### 2.2 Image Processing

The quantitative phase image of sperm cells was retrieved from the recorded four intensity images corresponding to 0, π/2, π and 3π/2 phase shift. The retrieved method can be found in previous reports^9^. The irradiance distribution at each point **r** = (*x, y*) in the image plane is a function of the phase shift, namely

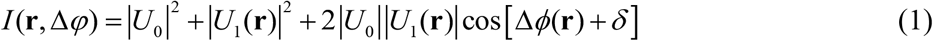

In Eq. 1, *U*_1_(**r**) is the scattered light from the sample and *U*_0_ is the incident light, *δ* is the phase shift introduced by the SLM. With the spatially varying phase shift, Δ*ϕ*(**r**) can be reconstructed as

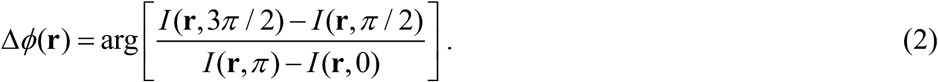

Then, the phase associated with the image field can be determined as^21^

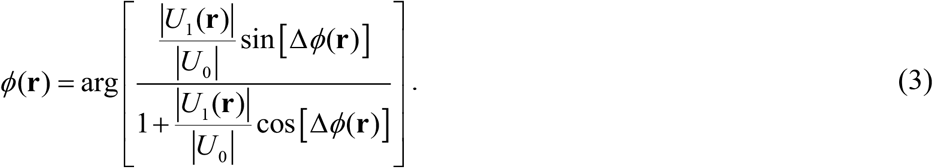

In order to obtain the thickness and refractive index each sperm cell was chosen as region of interest (ROI) from the quantitative phase image. For each group, the 100 ROIs were chosen and tiled into a mosaic using ImageJ. Then halo effect due to phase contrast illumination was removed for each selected ROI using a recent procedure developed in our laboratory^22^.

The halo removed ROIs were then further processed using ImageJ. First, the 30-pixel-wide segmented line selection width was carried out to select the sperm neck and tail. Second, the straighten plugin was applied to each new ROI to digitally remove the tail curvature. The 30-pixel values of each cross section along the straight line were retrieved and the profile along each tail was plotted.

## 3 Results

In this paper we measure, for the first time to our knowledge, both the topography and refractometer of sperm cells, in a spatially-resolved manner. We use a highly sensitive quantitative phase imaging instrument, CellVista SLIM Pro (Phi Optics, Inc.), which is described in Fig.1a. The SLIM module^23^ attaches to an existing phase contrast microscope and modulates the phase shift of the incident vs. scattered light in increment of π/2. To achieve this precise modulation, the objective pupil plane is relayed at the plane of a phase only Spatial Light Modulation (SLM). Without modulation on the SLM, we record a common phase contrast image. However, once a phase shift is applied on a mask that matches the ring of the objective ring, a different intensity image is obtained at the camera. Recording four intensity images corresponding to the 0, π/2, π and 3π/2 (Fig. 1b) phase shift, the phase map associated with the image field can be retrieved quantitatively and uniquely (Fig. 1c).

**Fig. 1.**
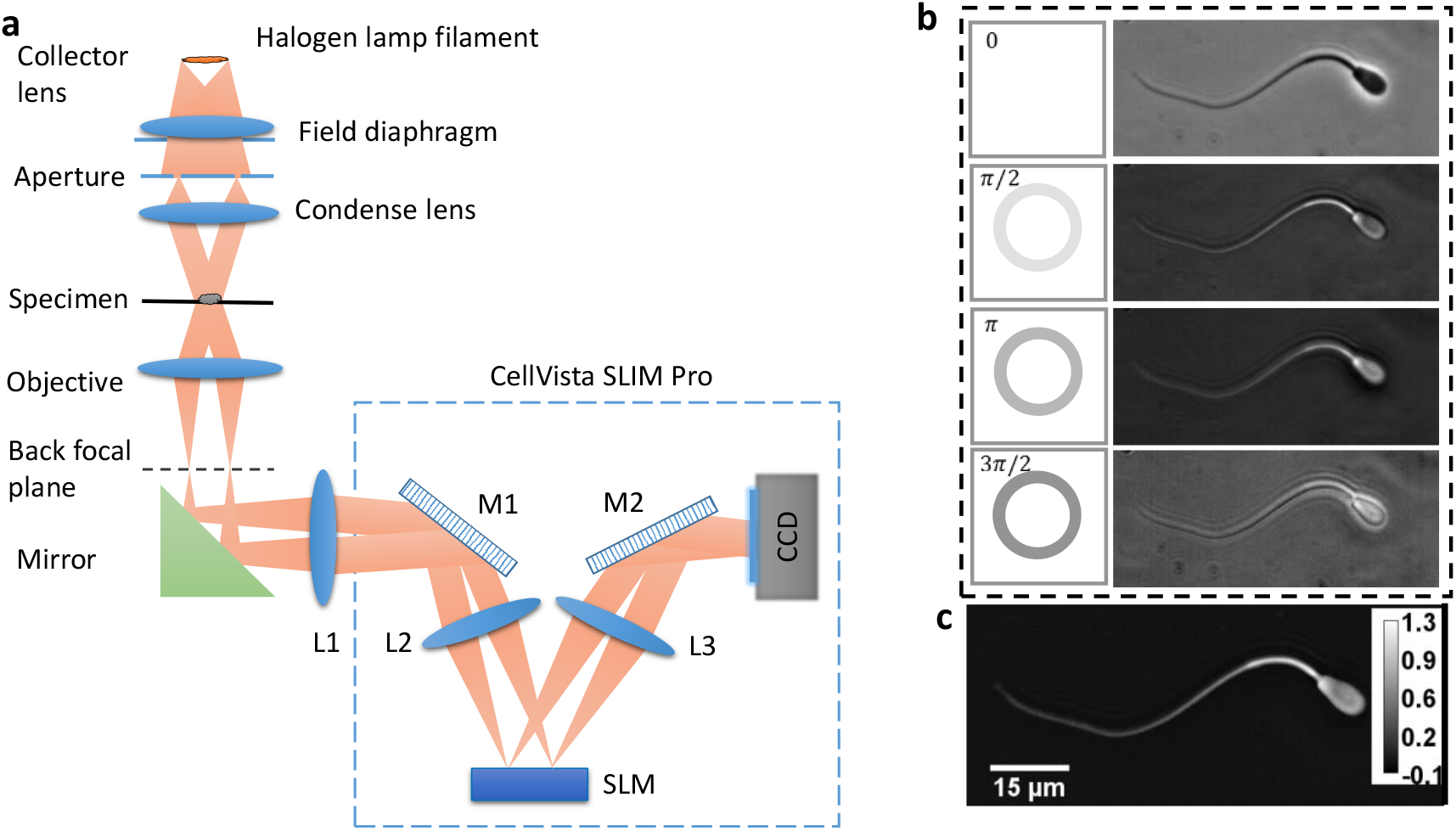
(a) Schematics of the spatial light interference microscope (SLIM). SLIM combines conventional phase contrast microscopy and a SLIM module (Cell Vista SLIM Pro, Phi Optics, Inc.). The SLIM module consists of a 4f lens system and a spatial light modulator (SLM) which produces phase modulation. (b) The phase rings and four intensity images corresponding to 0, π/2, π and 3π/2 phase shift recorded by the CCD. (c) High contrast SLIM quantitative-phase image created by combining the four intensity images. The color bar indicates phase in radian.

In order to decouple the cell thickness and refractive index from the optical path length information, we image the cells in two different media: air (n_0_=1) and PBS solution (n_1_=1.334). The cells were prepared as detailed in Materials and Methods. We imaged 100 cells for each of the air and PBS groups. As shown in Figs. 2–3, we notice immediately that the cell-induced phase shift is significantly lower in PBS vs. air, as expected for a smaller refractive index contrast. The spatial pathlength noise, evaluated as the standard deviation of a background area, shown by the green boxes in Figs. 2b–3b, is of the order of 1 nm in both cases (see the histograms insets). Note that this noise includes also irregularities in the background due to sample preparation, e.g., debris, solution in homogeneity, etc. Without a sample, the optical pathlength noise level achieved by SLIM is below one nanometer^6^.

**Fig. 2.**
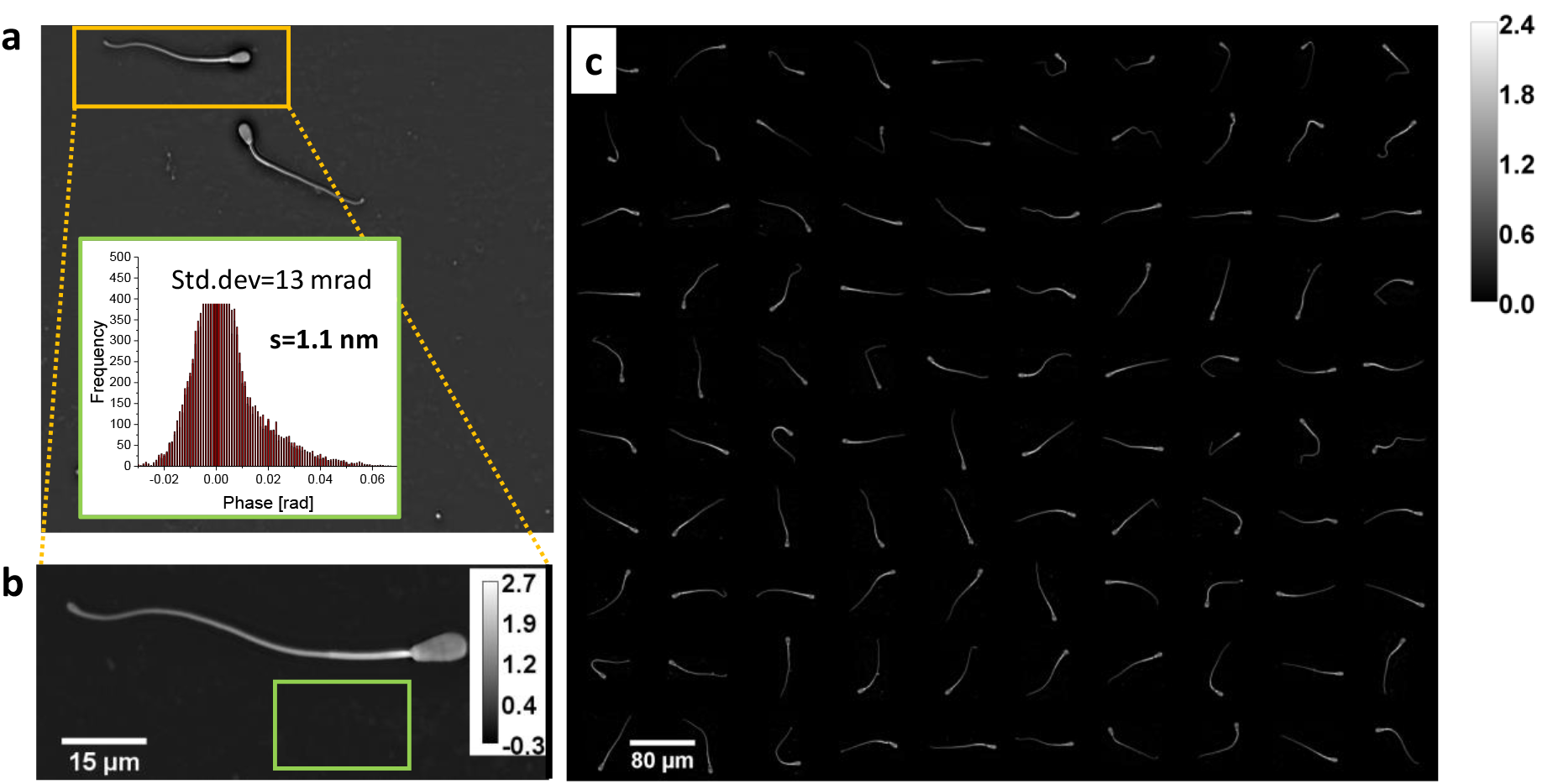
(a) Example of a single cell phase image. (b) The corresponding ROI of the “air” group. The standard deviation of a background area marked by the green boxes is shown in the histograms insets. (c) The montage of all the 100 cells measured from this group.

**Fig. 3.**
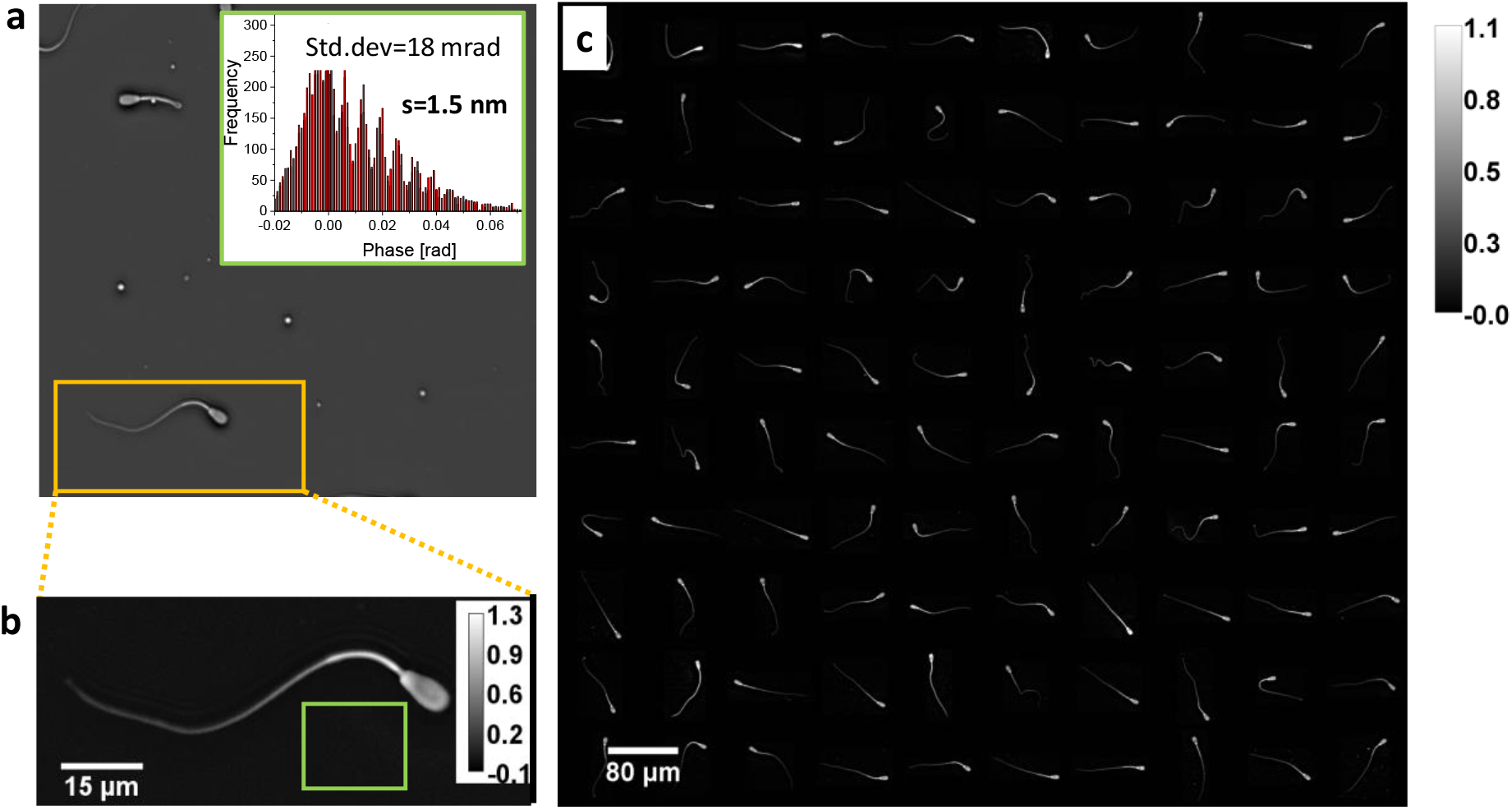
(a) Example of a single cell phase image. (b) The corresponding ROI of the “PBS” group. The standard deviation of a background area marked by the green boxes is shown in the histograms insets. (c) The montage of all the 100 cells measured from this group.

Each SLIM image provides morphological details that allow us to identify the hallmarks of the cell components (Fig. 4a). Interestingly, the nucleus of the cell appears to have a lower refractive index than the rest of the cell head, which seems to confirm the recent findings on eukaryotic cells^24, 25^. In order to perform averages across different cells, without artifacts due to the curvature, the tail of each cell was numerically “straighten” (Fig.4) using an ImageJ plugin developed by Meijering *et al*^26^. In essence, using this tool, we converted the 2D tail paths, into 1D trajectories. Once all tails are converted to lines, we align them such that the point where the neck ends is constant to all of them. With this procedure, we average the phase values of all the cells in each group. Fig. 5 shows the individual phase curves and the resulting two average profiles. The two-phase profiles depend on the average local refractive index, *n*(*x*), and thickness *h*(*x*), via the following system of equations

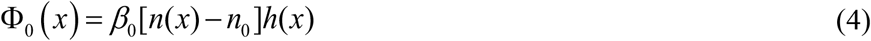

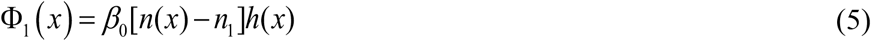

**Fig. 4.**
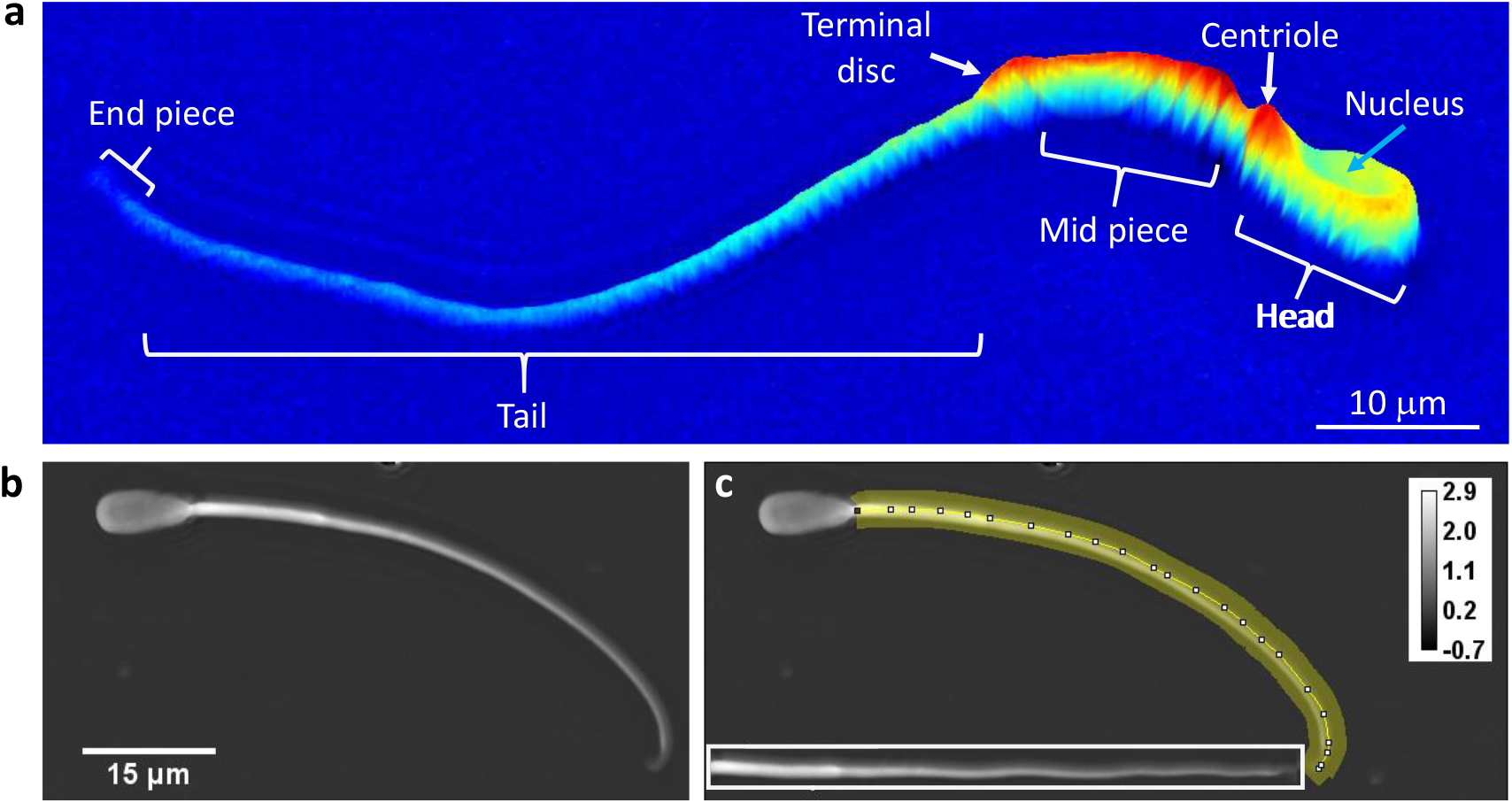
(a) A specific SLIM image with morphological details and hallmarks of the cellular components. (b) A single sperm with curved tail and (c) The corresponding 30-pixle-wide segmented line along the tail. The numerically straighten tail is shown in inset.

**Fig. 5.**
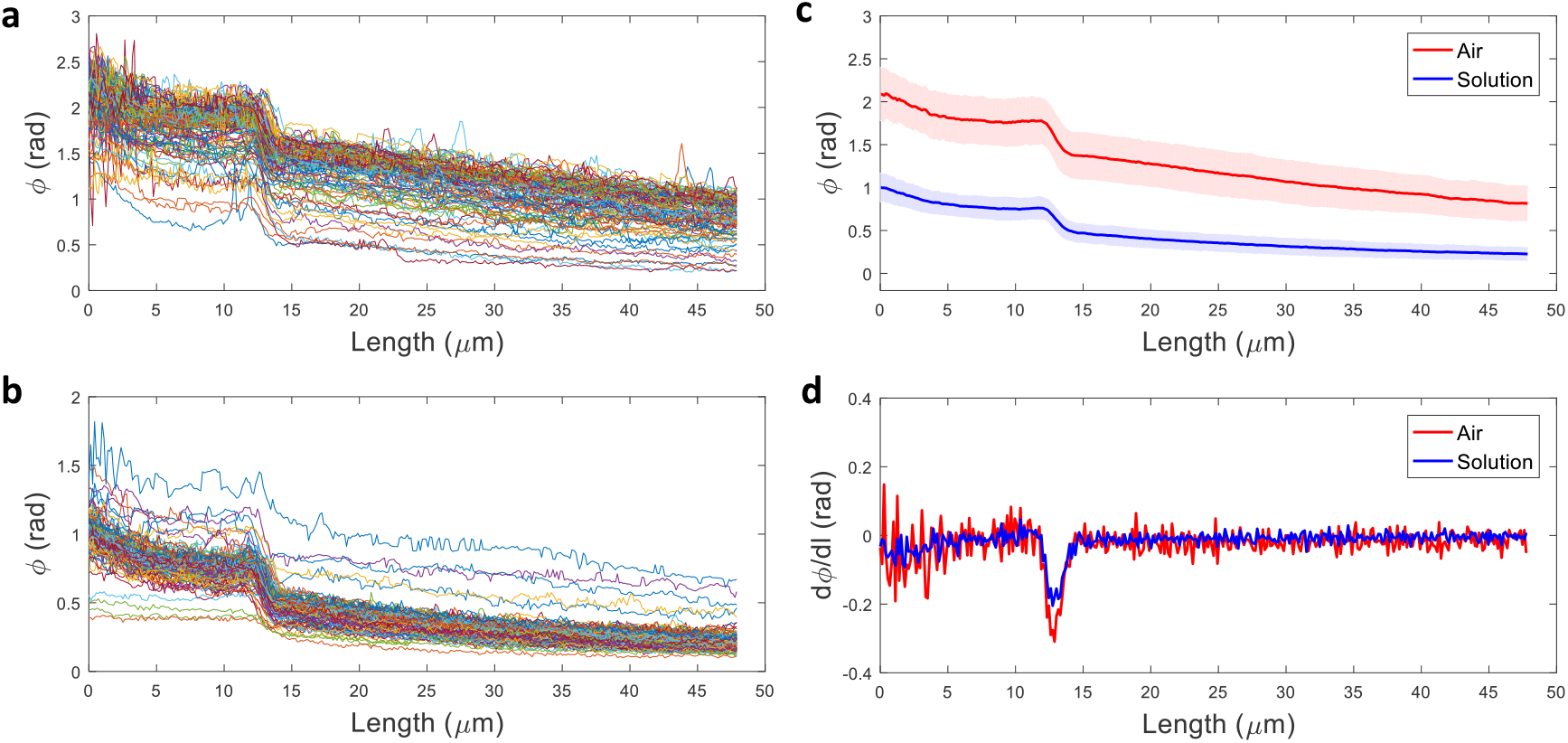
The individual phase curves of (a) the air group and (b) the PBS group. (c) The resulting two average profiles. (d) First-order derivative of the average profiles, which are used for aligning the thickness and refractive index profiles.

In Eqs 4–5, **Φ**_0_ and Φ_1_ are the phase shift in air and PBS, respectively, *β*_0_ is the wave number in vacuum *β*_0_ = 2*π*/*λ*, ***λ*** is the wavelength, and *x* is the coordinate along the tail. Solving the system of equations, we obtain

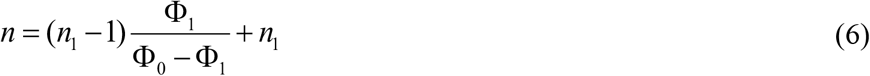

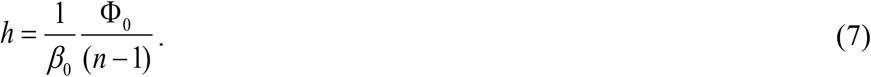

Note that the two phase measurements are characterized by a mean, **Φ**_0_ and Φ_1_, and standard deviation, *δ*Φ_0_ and *δ*Φ_1_, calculated over 100 cell measurements per group. As a result, the thickness and refractive index profiles are also characterized by standard deviations, ***δn*** and ***δh***, which are shown in Fig. 6.

**Fig. 6.**
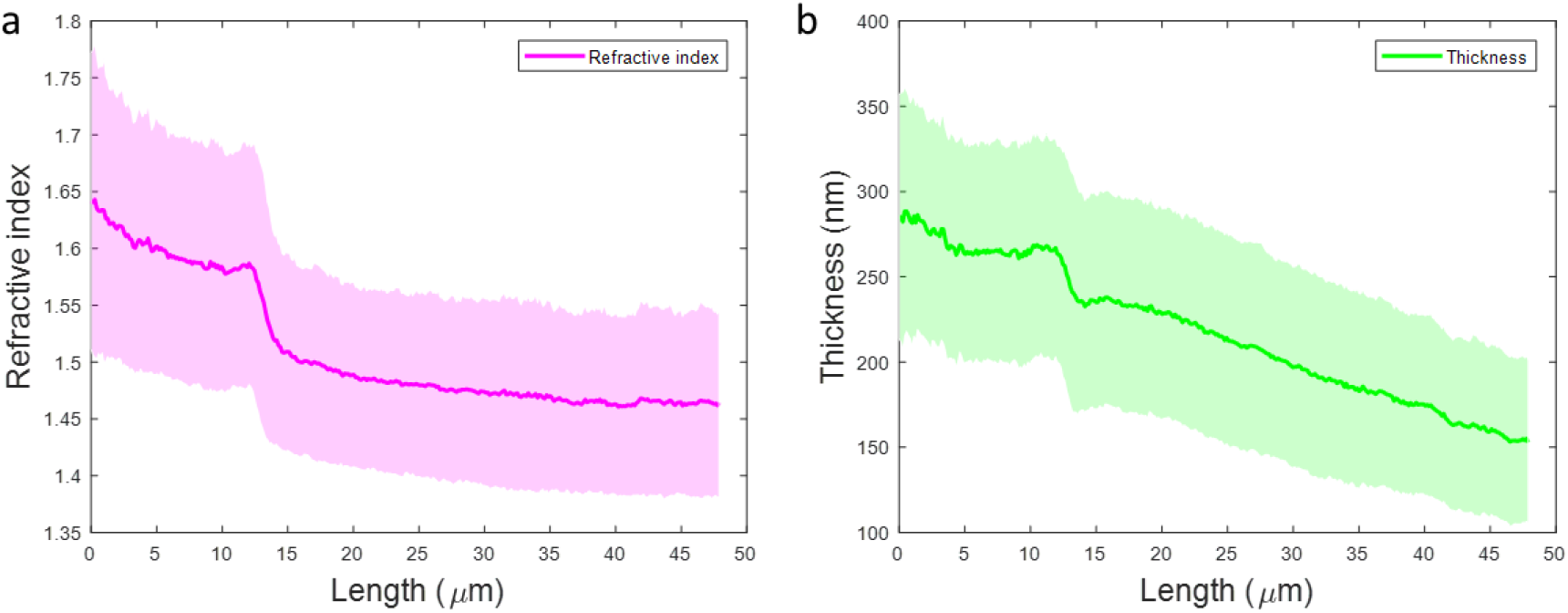
The thickness (a) and refractive index (b) profiles of spermatozoa. The thickness of the shaded area corresponds to the standard deviation.

The results show that the neck of the sperm is significantly different from the tail not just in terms of thickness, but also refractive index. Thus, the thickness decreases along the neck and tail from approximately 300 nm to 150 nm. Perhaps the most significant is that the refractive index of the neck reaches very high values, of 1.6 toward the head. The relatively high standard deviations include the intrinsic cell-to-cell variability, since they are the result of averages on two different cell assembles. The values of refractive index decrease toward the tail, down to 1.45, which is comparable to previous reported values. The refractive index of the neck is surprisingly high. It is well known that this region of the cell is packed with mitochondria^18^. These organelles are surrounded by lipid bilayers, which is likely responsible for such high values of refractive index.

## 4 Summary and Discussion

In summary, we measured for the first time to our knowledge simultaneous topography and refractometry of sperm cells. We performed imaging of cells suspended in two media of different refractive indices, which allowed us to retrieve separately both the thickness and refractive index of the cell tails. Using SLIM, a label-free method capable of providing nanoscale sensitivity to optical pathlength, we obtained detailed topography from single spermatozoa. Our technique was enabled by numerically eliminating tail curvatures. This procedure allowed us to average the results over a large number of cells in two measurement groups, characterized by different background refractive indices. Recent advances in SLIM technology allow us today to scan entire microscope slides and multi-well plates in a fully programmable and high-throughput manner^27^. The refractive index has been used before as a reporter for biomedical applications^28–31^. We anticipate that these new results will open a new area of applications by providing more accurate assays for IVF.

## Acknowledgments

This work was supported by the National Science Foundation (CBET-0939511 STC, DBI 14-50962 EAGER, IIP-1353368).

